# Selection of appropriate reference genes for quantitative real-time PCR in *Clerodendrum trichotomum*

**DOI:** 10.1101/625145

**Authors:** Yajie Hua, Yuanzheng Yue, Gongwei Chen, Taotao Yan, Wenjie Ding, Tingting Shi, Die Hu, Lianggui Wang, Xiulian Yang

## Abstract

*Clerodendrum trichotomum*, an important medicinal plant, has excellent salt tolerance and beautiful ornamental character. However, reliable reference genes for quantitative real-time PCR data (qRT-PCR) in *C. trichotomum* have not been investigated. Using our previous transcriptome data, 17 reference genes were selected in different tissues (leaves, flowers, fruits, stems, and roots) and under various abiotic stresses (salt, drought, flood, and heat) for *C. trichotomum*, using four different reference gene analysis software types: GeNorm, NormFinder, BestKeeper and ReFinder. The results identified *RPL*, *ACT* and *HSP70* as the three most suitable reference genes for tissues. Genes *ACT* and *AP-2* were most stably expressed under drought stress; *MDH* and *UBCE2* were stable under flooding stress; *RPL* and *UBCE2* were most stable under salt stress; and *MDH* and *EF-1A* were most appropriate under heat stress. For abiotic treatments, *RPL*, *MDH* and *AP-2* were the most stable reference genes; and *AP-2*, *RPL* and *ACT* were stably expressed in all examined samples. The expression profile of the genes for Na^+^/H^+^ Exchanger1 (*ClNHX1*) and laccase (*ClLAC*) were selected to validate the stability of the determined reference genes. Our study provided reliable normalization for gene expression analysis and ensured more accurate data for further molecular mechanism research in *C. trichotomum*.

## Introduction

*Clerodendrum*, a genus of flowering plants in the family Lamiaceae (Verbenaceae) (Wang et al., 2018), is widely distributed across China, Korea, Taiwan, Japan and India (Yamazaki., 1993). *Clerodendrum trichotomum* has been used for medicinal substances to act against human tumor cells (Wang et al., 2013a) and many new medicinal substances have been found in its tissues (Wang et al., 2013b; Xu et al., 2014). Moreover, *C. trichotomum* shows strong abiotic stress resistance and is easy to cultivate and manage (Mahmuc et al., 2008). As an ornamental plant, *C. trichotomum* has white petals, with a green calyx which turns red as the fruit ripens. The fruits (drupes) shift from white to bright blue and eventually dark blue in maturity. However, stable reference genes for *C. trichotomum* remain largely unknown and deeper research on molecular mechanisms in *C. trichotomum* is limited. Hence, selecting accurate reference genes for qRT-PCR analysis is urgently needed.

Generally, the reference genes used for plant research include *glyceraldehyde-3-phosphate dehydrogenase* (*GAPDH*), *Actin* (*ACT*), *elongation factor 1 alpha* (*EF-1A*), *elongation factor 4 alpha* (*EIF4a*), *ubiquitin* (*UBQ*), *Ubiquitin-conjugating enzyme E2* (*UBCE2*), *Tubulin* (*TUB*), *18S ribosomal RNA* (*18S*) and ribosomal protein (*RPL*) (Pfaffl et al., 2001; Lilly et al., 2011; Mamidala et al., 2011). With the development of transcriptome sequencing technology, more candidate genes have been identified (Machado et al., 2015). Stable reference genes have been confirmed in various plants, such as ornamental herbs (Fernandez et al., 2011; Qi et al., 2016), ornamental woody plants (Wang et al., 2014; Zhang et al., 2017), grasses (Yang et al., 2015; Takamori et al., 2017; Xu et al., 2017), fruits (Borges et al., 2014; Galli et al., 2015; Xu et al., 2015), vegetables (Jiang et al., 2014; Tian et al., 2015) and food crops (Hu et al., 2009; Rodriguez et al., 2016; Shivhare et al., 2016; Gines et al., 2017). Reference gene research has focused on plants with short growth cycles, with few studies performed on woody plants. However, woody plants in particular have higher biomass and longer growth period, and have a strong ability to deal with soil salinization and other problems (Zhang et al., 2017).

There has been no recent systematic screening of internal reference genes of *C. trichotomum*. Here, samples of *C. trichotomum* which were used for reference gene selection were collected from different provinces in China and named according to their place of sampling. Our previous physiological and molecular experiments showed that *C. trichotomum* sourced from Taian (TA) showed strong salt tolerance, and that sourced from Yancheng (YA) had the longest flowering period among all sources. Through the sequencing datasets of *C. trichotomum*, 17 reference genes were detected by qRT-PCR in roots, stems and different developmental stages of leaves, flowers and fruits: *ACT*, *PP2A*, *RPL*, *PK*, *18S*, *RAN*, *APT*, *SAND*, *PROF*, *MDH*, *EF-1A*, *UBCE2*, *AP-2*, *HSP70*, *TUA*, *UBQ* and *H3*. Expressions of these reference genes were analyzed in different tissues and various abiotic treatments. Then, four software types were used to calculate the qRT-PCR results: BestKeeper (Pfaffl et al., 2004), GeNorm (Vandesompele et al., 2002), NormFinder (Andersen et al., 2004) and ReFinder (http://150.216.56.64/referencegene.php). These indicated that *AP-2*, *RPL* and *ACT* were the most suitable genes in all samples.

The Na^+^/H^+^ Exchanger1 gene (*ClNHX1*) had been identified as crucial in strengthening the different abiotic stress tolerances of plants, especially salt tolerance (Jha et al., 2011; Sahoo et al., 2016), as has the gene for laccase (*ClLAC*) which is involved in lignin synthesis and is expressed in different organs (Kitajima et al., 2017; Li et al., 2018). In this study, *ClNHX1* and *ClLAC* of *C. trichotomum* were selected to validate the reliability of target reference genes, and the results indicated that our reference genes selection was reliable. This work provided a solid reference gene resource for further molecular research of *C. trichotomum*.

## Materials and methods

### Plant materials and stress treatments

The *C. trichotomum* samples from TA and YA were used in this study. Plants were grown in the Baima Research and Teaching Base of Nanjing Forestry University (Nanjing, China). Five tissue samples were collected: stems, roots, leaves (young, mature and old leaves), flowers (flower bud, early flowering and full flowering stages) and fruits (early, mid and late development). All materials were immediately frozen in liquid nitrogen and then kept at −80°C until used.

Cuttings were planted in pots filled with a 1:1:1:2 mixture of perlite: vermiculite:peat:sand. Plants were maintained in a growth chamber (RDN-1000-3, Southeast Instrument Factory, Ningbo, China) with 14-h photoperiod, 25/21°C (day/night), light intensity 180 mmol/m^2^/s and relative humidity of 60% for 3 months. Cuttings were put into quarter-strength Hoagland medium (Hoagland et al., 1905) adapted for one week before performing abiotic stress treatments. For drought and salinity stress treatments, the cuttings were transferred into nutrient solution with 100 mmol/l NaCl or 15% PEG6000, respectively. For flooding treatments, plants were maintained in rainwater. The plants were preserved in the growth chamber at 42/35°C (day/night) for heat stress treatment. The leaves of three plants per replicate were collected following salinity, drought and flood stress treatments at 0, 2, 6, 12, 24, 48 and 72 h. For thermal stress treatment, the leaves of three plants per replicate were collected at 0, 2, 6, 12, 24 and 48 h. Then the leaves were immediately frozen in liquid nitrogen and stored at −80°C until used.

### Total RNA isolation and cDNA synthesis

Total RNA was isolated using EASY spin Plus (Aidlab Biotechnologies Co. Ltd. Beijing, China), and the integrity of total RNA was confirmed by 2.0% (w/v) agarose gel electrophoresis. Then the quality of RNA (A260/280; A260/230) were measured with the UV5NANO (Mettler Toledo, Switzerland). cDNA synthesis was carried out using 2 μg of total RNA in a final volume of 20 μl by EasyScript One-Step gDNA Removal and cDNA Synthesis SuperMix (TransGen Biotech Inc. Beijing, China), finally, the cDNA was diluted 10-fold with nuclease-free water for qRT-PCR (Yue et al., 2018).

### Selection of reference gene sequences and primer design

Candidate genes were selected from the *C. trichotomum* transcriptome using the Illumina Hi-Seq™ 3000 platform. A total of 17 candidate reference genes were identified by Blast (https://blast.ncbi.nlm.nih.gov/Blast.cgi): *ACT*, *PP2A*, *RPL*, *PK*, *18S*, *RAN*, *APT*, *SAND*, *PROF*, *MDH*, *EF-1A*, *UBCE2*, *AP-2*, *HSP70*, *TUA*, *UBQ* and *H3*. Primers were designed by Primer Premier 5 software and based on our usual methods (Mu et al., 2017) (Table 1). In order to detect the specificity of the amplicon, the amplified products of each gene were verified by 2.0% (W/V) agarose gel electrophoresis. The qRT-PCR amplification reactions were performed by using the Applied Biosystems StepOne PCR System (Thermo Fisher Scientific, USA) and SYBR Premix Ex Taq™ (TakaRa, Japan). The reaction volume was 10μl, in which the cDNA template was 1 μl, the ROX Reference Dye II was 0.2 μl, the forward and reverse amplification primers were 0.4 μl, the 2× SYBR Primer Ex Taq™ was 5 μl, and the ddH_2_O was 3 μl. The reaction procedure was using our usual methods(Yang et al., 2018; Yue et al., 2017). All qRT-PCR reactions were repeated three times biologically and three times technically, and three times template-free negative control.Ultimately, the reliability of the target reference genes were verified by using *ClNHX1* and *ClLAC* genes, the reaction volume and procedure were the same as above.

**Table 1.**
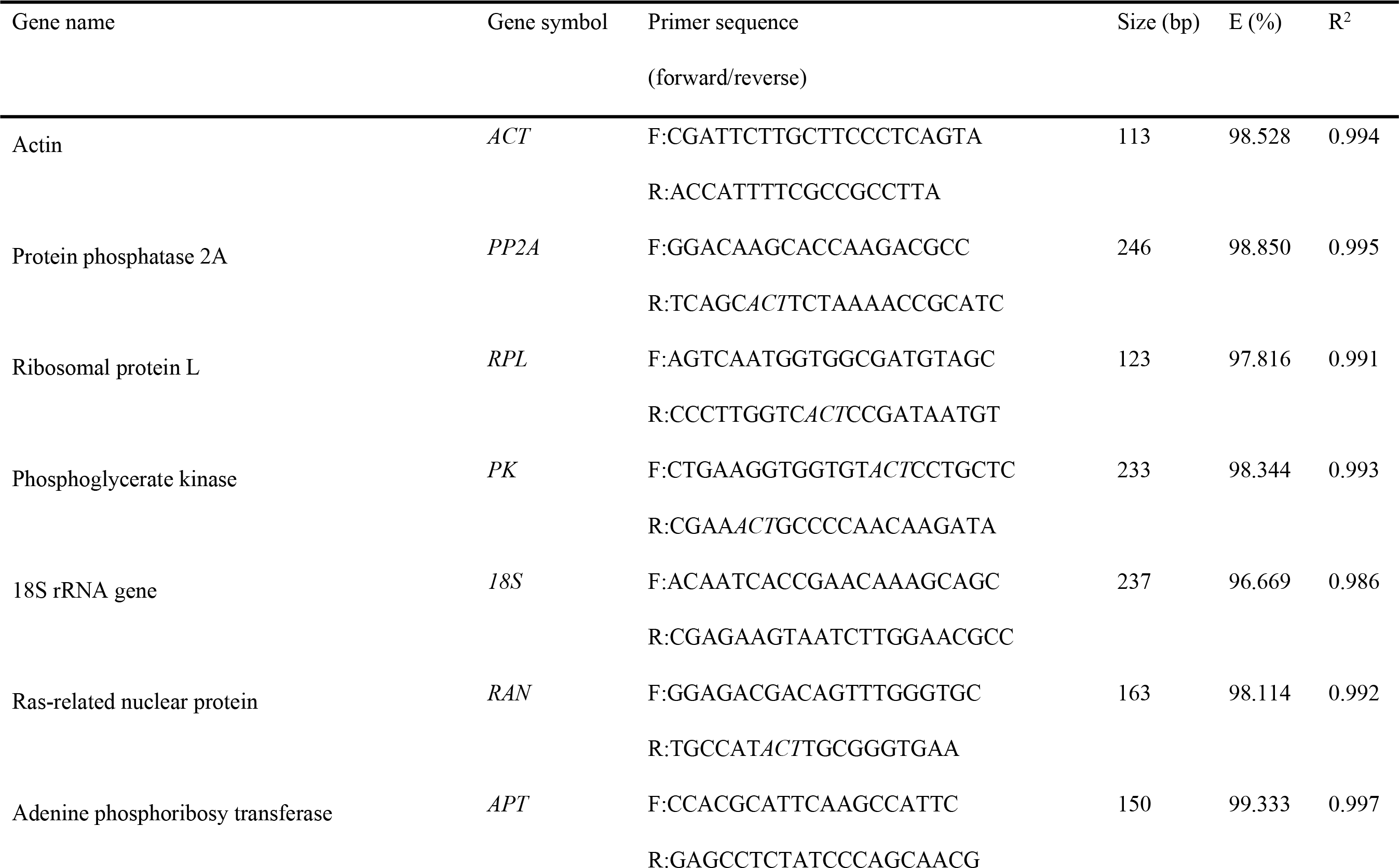
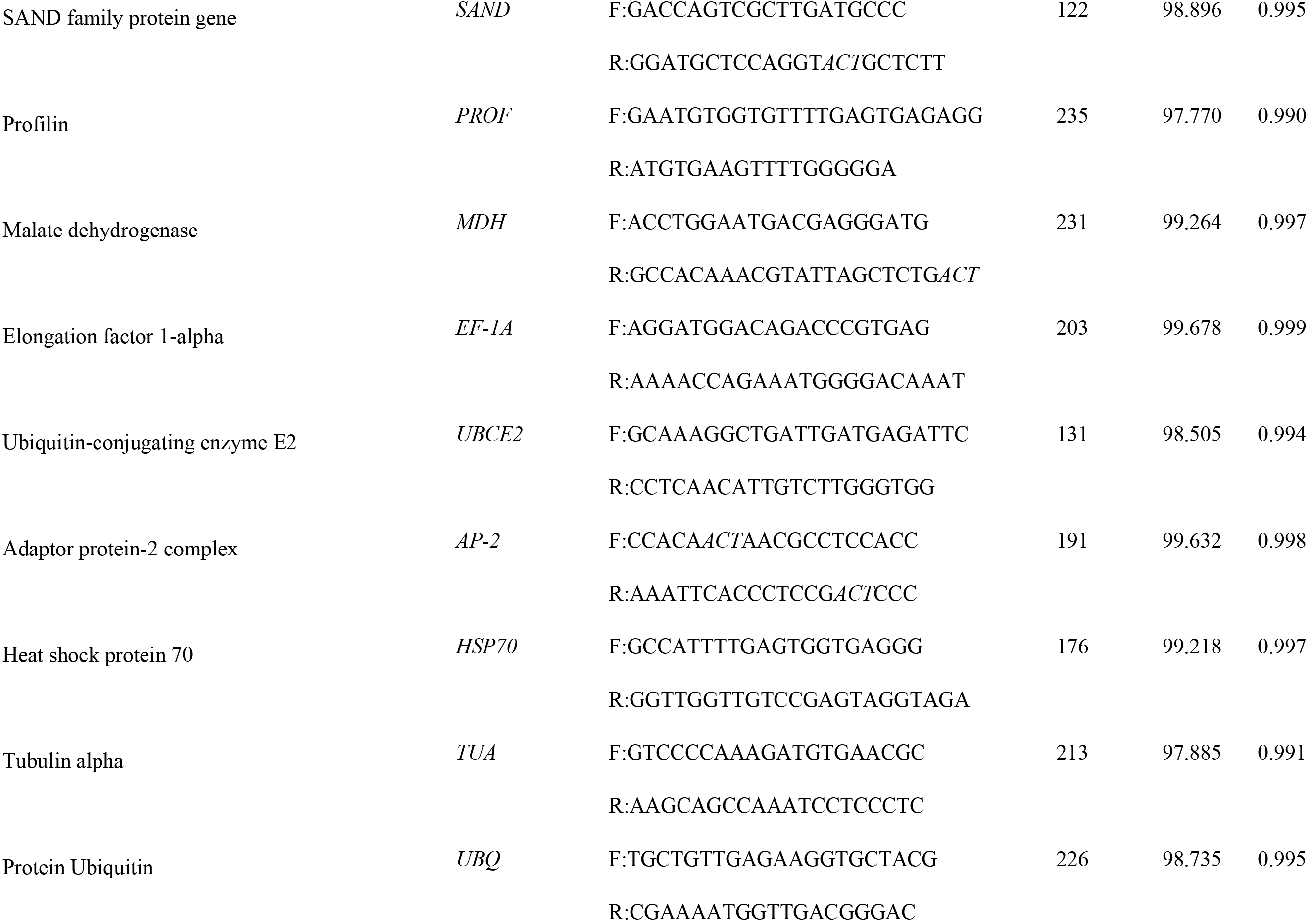
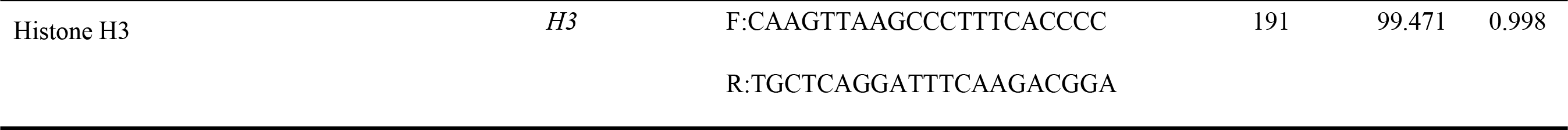
Genes and primer sets used for qRT-PCR.

### Calculation of PCR efficiencies

Five-fold series of dilutions (1, 1/5, 1/25, 1/125 and 1/625) were performed on the cDNAs including the equimolar quantities of all samples to generate standard curves. The amplification efficiency (E) : E (%) = (10^−1/slope^ − 1) × 100 for each reference gene was calculated, and the slope was the standard curve slope calculated by Applied Biosystems StepOne (Thermo Fisher Scientific) (Savard P et al., 2009).

### Statistical analysis of gene expression

NormFinder, GeNorm and BestKeeper were used to select suitable reference genes, and ReFinder (http://150.216.56.64/referencegene.php) used for sequencing the stability of these reference genes. When using GeNorm, the Δct value was first calculated—the smallest ct value in the samples was subtracted from the ct value of other samples to get the Δct value—using Excel to calculate the corresponding gene of 2^−Δct^ and import the 2^−Δct^ value into GeNorm software for analysis (Schmittgen et al., 2008).

## Results

### Assessment of primer specificity and PCR amplification efficiency

The single peak on the melting curve of each reference gene represented the specification of each gene (Fig. 1). The amplification efficiency of theses 17 genes ranged from 96.67% (*18S*) to 99.68% (*EF-1A*) (Fig. 1), indicating that these genes had reliable amplification efficiency and were suitable for further gene expression experiments. The correlation coefficients (R^2^) of the standard curve ranged from 0.986(*18S*) to 0.999 (*EF-1A*) (Table 1). The level of gene expression was determined by cycle threshold (CT), and the CT values of 17 reference genes ranged from 19.66 (*ACT*) to 28.35 (*PROF*). These candidate genes showed different levels of expression, *ACT* expression was the highest and *PROF* expression was the lowest (Figure 2).

**Fig. 1.**
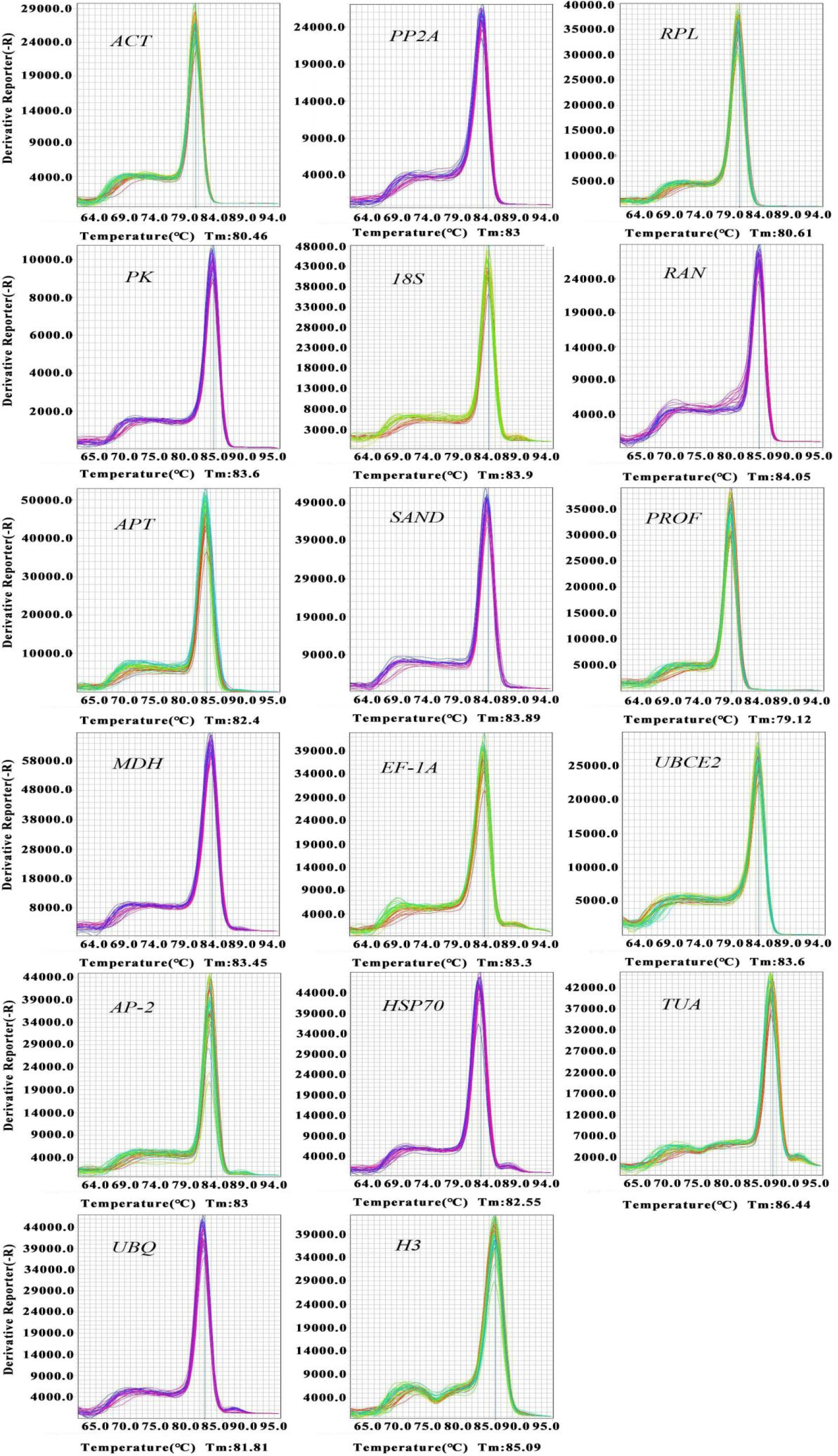
Primer specificity. Melting curves of 17 reference genes (lines 1–17: *ACT*, *PP2A*, *RPL*, *PK*, *18S*, *RAN*, *APT*, *SAND*, *PROF*, *MDH*, *EF-1A*, *UBCE2*, *AP-2*, *HSP70*, *TUA*, *UBQ* and *H3*) show simple peaks.

**Fig. 2.**
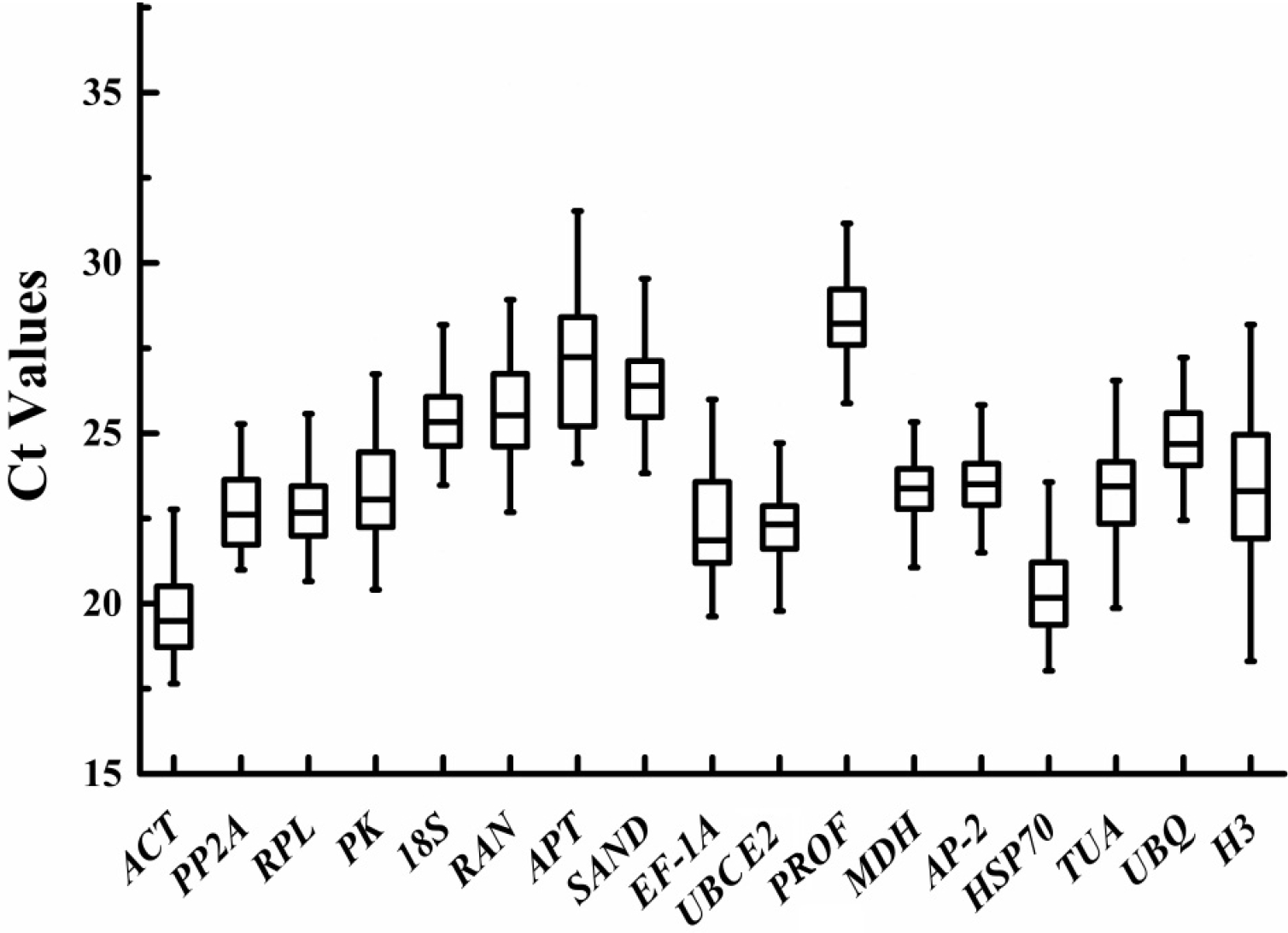
Ct values of 17 candidate reference genes in all samples. A line on the box depicts the middle belt. The upper and lower boxes represent the 25th and 75th percentiles, respectively. The whiskers show maximum and minimum values.

### GeNorm analysis

Genorm software was used to analyze the expression stability values (M) of different samples to determine the most stable genes. The gene with the lowest M value was considered to be the most stable and vice versa; recommended stable expression of M-value is < 1.5 (Zhang et al. 2019; Vandesompele et al. 2002). GeNorm demonstrated that *RPL* and *HSP70* were the best genes in TA tissues and all tissues (both TA and YA source); but in YA tissues, *RAN* and *HSP70* were more stable. Under drought stress, *ACT* and *PP2A* expressed most stably in TA source, and *ACT* and *AP-2* showed most stable expreesion in YA source. For flooding stress, *UBCE2* and *MDH* were most stable in TA and for YA, *PP2A* and *AP-2* were the most stable. The most stable genes under salt stress for both TA and YA were *RPL* and *AP-2*. For heat stress, *EF-1A* and *MDH* were most suitable for TA source, and *EF-1A* and *UBCE2* were most suitable for YA. For all abiotic stress treatments, *MDH* and *AP-2* were the most stable (i.e. lowest M-value). In the context of the total sample set, *ACT* and *AP-2* were the most stable. Genes *ACT* and *PP2A* were the most stable in all TA samples, and *RPL* and *AP-2* were the most stable in YA samples. Gene *H3* was always ranked as unstable (Table 2).

**Table 2.** Ranking order of candidate genes calculated by GeNorm, NormFinder, BestKeeper. (TA, Taian; YA, Yancheng)..

| Rank | GeNorm |  |  |  |  |  | NormFinder |  |  |  |  |  | BestKeeper |  |  |  |  |  |
| --- | --- | --- | --- | --- | --- | --- | --- | --- | --- | --- | --- | --- | --- | --- | --- | --- | --- | --- |
|  | Best |  |  | Least |  |  | Best |  |  | Least |  |  | Best |  |  | Least |  |  |
|  | 1 | 2 | 3 | 1 | 2 | 3 | 1 | 2 | 3 | 1 | 2 | 3 | 1 | 2 | 3 | 1 | 2 | 3 |
| TA tissues | *RPL* | *HSP70* | *UBCE2* | *TUA* | *APT* | *H3* | *RPL* | *ACT* | *HSP70* | *TUA* | *APT* | *H3* | *PP2A* | *RAN* | *AP-2* | *APT* | *TUA* | *H3* |
| YA tissues | *RAN* | *HSP70* | *RPL* | *PK* | *TUA* | *H3* | *RPL* | *HSP70* | *AP-2* | *TUA* | *PK* | *H3* | *AP-2* | *SAND* | *PP2A* | *H3* | *MDH* | *TUA* |
| All tissues | *RPL* | *HSP70* | *RAN* | *TUA* | *APT* | *H3* | *RPL* | *ACT* | *RAN* | *PK* | *APT* | *H3* | *AP-2* | *18S* | *ACT* | *APT* | *TUA* | *H3* |
| TA drought | *ACT* | *PP2A* | *EF-1A* | *18S* | *H3* | *PROF* | *AP-2* | *APT* | *RAN* | *18S* | *H3* | *PROF* | *ACT* | *PP2A* | *AP-2* | *PROF* | *TUA* | *H3* |
| YA drought | *ACT* | *AP-2* | *UBCE2* | *EF-1A* | *PROF* | *H3* | *AP-2* | *MDH* | *UBCE2* | *PROF* | *EF-1A* | *H3* | *ACT* | *UBCE2* | *AP-2* | *PK* | *PROF* | *EF-1A* |
| TA flood | *UBCE2* | *MDH* | *PK* | *H3* | *APT* | *RAN* | *MDH* | *PK* | *PP2A* | *H3* | *APT* | *RAN* | *PP2A* | *MDH* | *UBCE2* | *EF-1A* | *APT* | *RAN* |
| YA flood | *PP2A* | *AP-2* | *UBQ* | *APT* | *PROF* | *H3* | *MDH* | *UBCE2* | *EF-1A* | *APT* | *PROF* | *H3* | *MDH* | *EF-1A* | *TUA* | *SAND* | *APT* | *H3* |
| TA salt | *RPL* | *AP-2* | *UBCE2* | *PK* | *APT* | *H3* | *MDH* | *UBCE2* | *RPL* | *EF-1A* | *APT* | *H3* | *ACT* | *UBCE2* | *RAN* | *SAND* | *APT* | *H3* |
| YA salt | *RPL* | *AP-2* | *UBCE2* | *H3* | *EF-1A* | *18S* | *RPL* | *UBCE2* | *AP-2* | *H3* | *EF-1A* | *18S* | *RPL* | *PROF* | *UBQ* | *APT* | *18S* | *EF-1A* |
| TA heat | *EF-1A* | *MDH* | *SAND* | *18S* | *APT* | *RAN* | *MDH* | *RPL* | *SAND* | *18S* | *APT* | *RAN* | *AP-2* | *PROF* | *UBQ* | *18S* | *APT* | *RAN* |
| YA heat | *EF-1A* | *UBCE2* | *MDH* | *PROF* | *UBQ* | *H3* | *MDH* | *AP-2* | *EF-1A* | *HSP70* | *UBQ* | *H3* | *RPL* | *PROF* | *UBCE2* | *HSP70* | *RAN* | *18S* |
| All abiotic stress | *MDH* | *AP-2* | *RPL* | *RAN* | *APT* | *H3* | *RPL* | *UBCE2* | *SAND* | *APT* | *RAN* | *H3* | *MDH* | *AP-2* | *UBQ* | *RAN* | *APT* | *H3* |
| TA all samples | *ACT* | *PP2A* | *AP-2* | *RAN* | *APT* | *H3* | *RPL* | *ACT* | *SAND* | *RAN* | *APT* | *H3* | *AP-2* | *MDH* | *PP2A* | *RAN* | *APT* | *H3* |
| YA all samples | *RPL* | *AP-2* | *ACT* | *EF-1A* | *PK* | *H3* | *AP-2* | *RPL* | *ACT* | *APT* | *PK* | *H3* | *AP-2* | *SAND* | *RPL* | *EF-1A* | *PK* | *H3* |
| Total samples | *ACT* | *AP-2* | *PP2A* | *RAN* | *APT* | *H3* | *RPL* | *AP-2* | *SAND* | *PROF* | *APT* | *H3* | *AP-2* | *MDH* | *SAND* | *RAN* | *APT* | *H3* |

The paired variation V-value of the normalized factor was calculated after a new internal reference gene was introduced. Then, the most suitable number of internal reference genes was determined; if Vn/(Vn + 1) < 0.15, then the number of reference genes needed was n, otherwise, the number needed was n + 1 (Vandesompele et al., 2002). For abiotic stresses including drought, flood, salt and heat stress, the pairwise variation of V2/3 was < 0.15, indicating that two reference genes were sufficient. For all abiotic stress samples, V5/6 < 0.15. In TA tissues, V3/4 < 0.15, which meant that three reference genes should be used as internal controls. Four reference genes were selected according to V3/4 values of YA tissues. In all TA samples, V5/6 < 0.15 and in all YA samples V5/6 was close to 0.15. In total samples, because V5/6 was close to 0.15, five reference genes would be sufficient for normalizing the samples (Fig. 3)

**Fig. 3.**
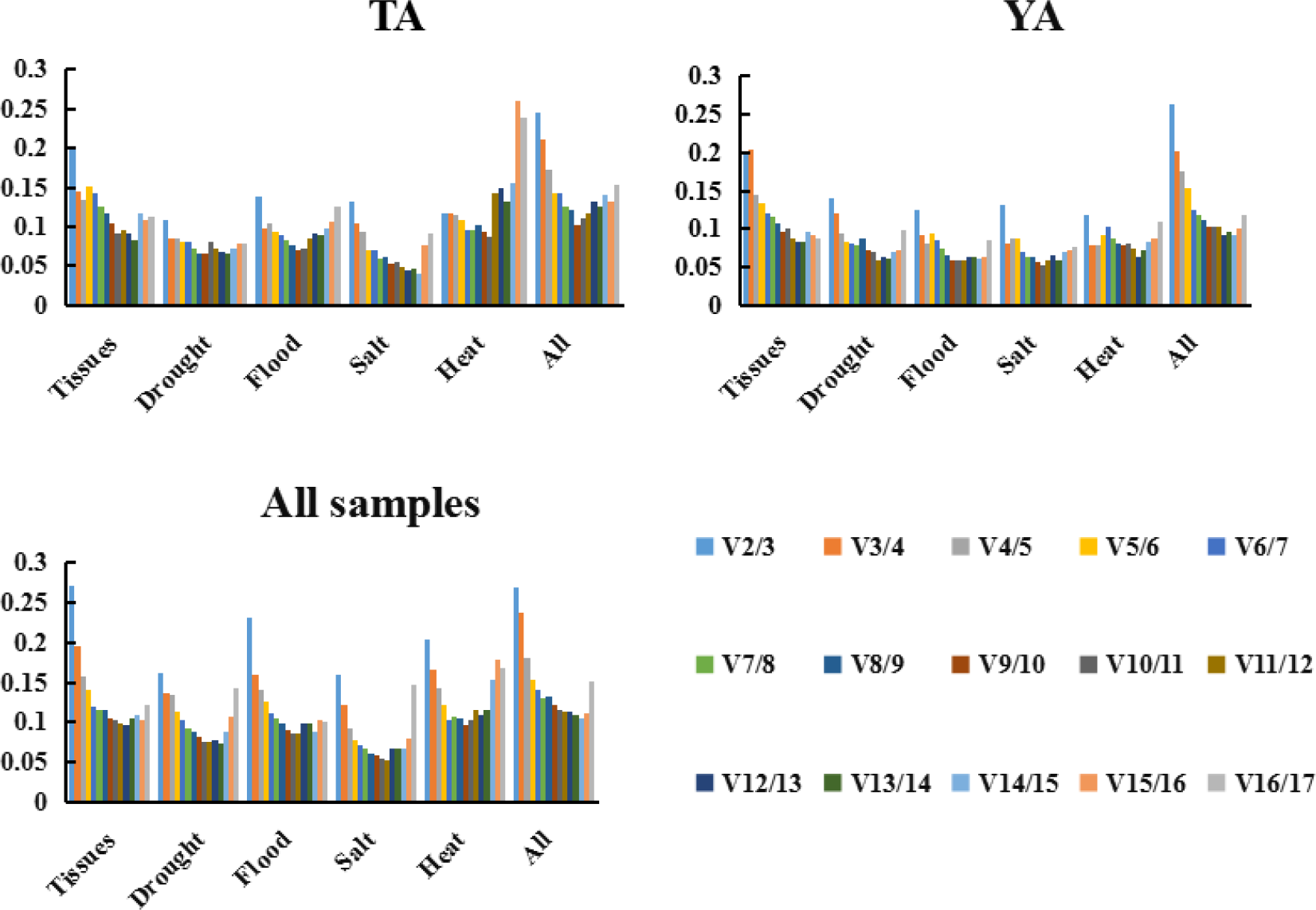
Pairwise variation (V-value) of reference genes calculated by GeNorm. The Vn/(Vn + 1) values were used to analyze the optimal numbers of reference genes.

### NormFinder analysis

NormFinder selected the reference gene with the lowest stable expression value. NormFinder only screened out one suitable internal reference gene (Andersen et al., 2004). For all tissue samples, *RPL* was the best gene (both in TA and YA tissues). In the subset of TA and YA under drought stress, *AP-2* was the most stable. For flood stress, *MDH* was the most stable in TA and YA. For salt stress, *MDH* was the best gene followed by *UBCE2* and *RPL* in TA; however, in YA, *RPL* was the most stable, followed by *UBCE2* and *AP-2*. For heat stress, *MDH* was the most stable gene for both TA and YA. For all abiotic stress samples, *RPL* was the most stable. In all TA samples, *RPL* was the most stable gene; for YA, *AP-2* was the most stable followed by *RPL*. In total samples, *AP-2* was the most stable followed by *RPL*. The *H3* was unstably expressed in all tissue and stress samples (Table 2).

### BestKeeper analysis

BestKeeper calculated the most relevant genes by analyzing the correlation coefficient (r), standard deviation (SD) and coefficient of variation (CV) between each gene and then comparing the individual values. Genes with larger r and smaller SD were considered more stable. Expression of reference gene > 1 was considered unsuitable for internal control. BestKeeper only filtered out one of the most suitable internal reference genes (Pfaffl et al., 2004). The *PP2A* was the most stable gene for TA tissues and *AP-2* was the most stable for YA tissues and all tissues; *ACT* was suitable for drought stress and *MDH* for heat stress. For flood stress, *PP2A* had the minimum CV ± SD followed by *MDH* in TA; however, in YA, *MDH* was the most stable. In the salt-stress experiment, *ACT* was the most stable gene in TA and *RPL* the most stable in YA. The *MDH* was the most stable for all abiotic stresses. For TA samples, YA samples and all samples, *AP-2* was the most suitable gene as an internal control (Table 2).

### Comprehensive stability analysis of reference genes

The best reference genes were combined by ReFinder, which showed that *RPL*, *HSP70* and *ACT* were suitable for TA tissues, and *RPL*, *HSP70* and *AP-2* were suitable for YA tissues. Among all tissue samples, *RPL*, *ACT* and *HSP70* were the best reference genes. The *ACT* and *AP-2* were suitable for drought stress, *MDH* and *UBCE2* for flood stress, *RPL* and *UBCE2* for salt stress, *MDH* and *EF-1A* for heat stress and *RPL*, *MDH* and *AP-2* for all abiotic stress treatments. The three best reference genes for all TA samples were *RPL*, *ACT* and *PP2A* and for all YA samples, *AP-2*, *RPL* and *ACT* were more stable. Considering all experiments, *AP-2*, *RPL* and *ACT* were the most suited as reference genes in *C*. *trichotomum* (Table 3).

**Table 3.** Stability of reference gene expression as recommended comprehensive ranking calculated by ReFinder (TA, Taian; YA, Yancheng).

| Method | 1 | 2 | 3 | 4 | 5 | 6 | 7 | 8 | 9 | 10 | 11 | 12 | 13 | 14 | 15 | 16 | 17 |
| --- | --- | --- | --- | --- | --- | --- | --- | --- | --- | --- | --- | --- | --- | --- | --- | --- | --- |
| TA tissues | *RPL* | *HSP70* | *ACT* | *AP-2* | *PP2A* | *UBCE2* | *RAN* | *MDH* | *18S* | *PK* | *UBQ* | *EF-1A* | *SAND* | *PROF* | *TUA* | *APT* | *H3* |
| YA tissues | *RPL* | *HSP70* | *AP-2* | *RAN* | *ACT* | *EF-1A* | *PP2A* | *SAND* | *UBCE2* | *APT* | *18S* | *MDH* | *PROF* | *PK* | *UBQ* | *TUA* | *H3* |
| All tissues | *RPL* | *ACT* | *HSP70* | *AP-2* | *RAN* | *SAND* | *UBCE2* | *18S* | *EF-1A* | *PP2A* | *UBQ* | *MDH* | *PROF* | *PK* | *TUA* | *APT* | *H3* |
| TA drought | *ACT* | *AP-2* | *PP2A* | *APT* | *EF-1A* | *RAN* | *UBCE2* | *SAND* | *MDH* | *HSP70* | *TUA* | *PK* | *RPL* | *18S* | *UBQ* | *H3* | *PROF* |
| YA drought | *AP-2* | *ACT* | *UBCE2* | *MDH* | *RAN* | *UBQ* | *PP2A* | *HSP70* | *APT* | *SAND* | *RPL* | *PK* | *18S* | *TUA* | *H3* | *PROF* | *EF-1A* |
| TA flood | *MDH* | *PP2A* | *UBCE2* | *PK* | *AP-2* | *TUA* | *SAND* | *UBQ* | *ACT* | *RPL* | *HSP70* | *PROF* | *EF-1A* | *18S* | *H3* | *APT* | *RAN* |
| YA flood | *MDH* | *UBCE2* | *EF-1A* | *AP-2* | *UBQ* | *TUA* | *PP2A* | *PK* | *18S* | *ACT* | *HSP70* | *RPL* | *RAN* | *SAND* | *PROF* | *APT* | *H3* |
| TA salt | *UBCE2* | *RPL* | *AP-2* | *MDH* | *RAN* | *ACT* | *UBQ* | *18S* | *TUA* | *PP2A* | *PROF* | *EF-1A* | *HSP70* | *PK* | *SAND* | *APT* | *H3* |
| YA salt | *RPL* | *UBCE2* | *AP-2* | *MDH* | *ACT* | *UBQ* | *PP2A* | *PROF* | *RAN* | *PK* | *TUA* | *SAND* | *HSP70* | *APT* | *H3* | *EF-1A* | *18S* |
| TA heat | *MDH* | *EF-1A* | *SAND* | *RPL* | *ACT* | *AP-2* | *UBCE2* | *PK* | *TUA* | *PROF* | *UBQ* | *PP2A* | *HSP70* | *H3* | *18S* | *APT* | *RAN* |
| YA heat | *MDH* | *EF-1A* | *UBCE2* | *AP-2* | *RPL* | *SAND* | *ACT* | *TUA* | *PROF* | *PP2A* | *PK* | *APT* | *RAN* | *UBQ* | *18S* | *HSP70* | *H3* |
| All abiotic | *RPL* | *MDH* | *AP-2* | *UBCE2* | *SAND* | *TUA* | *ACT* | *UBQ* | *PP2A* | *PK* | *HSP70* | *EF-1A* | *PROF* | *18S* | *RAN* | *APT* | *H3* |
| TA all samples | *RPL* | *ACT* | *PP2A* | *AP-2* | *SAND* | *MDH* | *UBCE2* | *PK* | *HSP70* | *UBQ* | *EF-1A* | *TUA* | *18S* | *PROF* | *RAN* | *APT* | *H3* |
| YA all samples | *AP-2* | *RPL* | *ACT* | *SAND* | *MDH* | *RAN* | *UBCE2* | *PP2A* | *HSP70* | *TUA* | *18S* | *UBQ* | *PROF* | *EF-1A* | *APT* | *PK* | *H3* |
| Total | *AP-2* | *RPL* | *ACT* | *SAND* | *MDH* | *UBCE2* | *PP2A* | *HSP70* | *UBQ* | *EF-1A* | *TUA* | *18S* | *PK* | *PROF* | *RAN* | *APT* | *H3* |

### Reference gene validation

To validate the identified reference genes, qRT-PCR was applied to evaluate expression of *ClNHX1* under different abiotic stress experimental conditions and of *ClLAC* in different tissues. Expressions of *ClNHX1* and *ClLAC* were normalized using the most stable combination of reference genes and both single compared with the least stable reference gene in each stress subset (Fig. 4). For drought stress, *ClNHX1* expression increased at 2 and 24 h, and decreased at other times. For salt stress, *ClNHX1* was upregulated during 0–48 h and downregulated thereafter. For flooding stress, *ClNHX1* was upregulated during 0–6 h and downregulated thereafter. For heat stress, *ClNHX1* expression was upregulated at 2 and 12 h and downregulated at other times. The *ClLAC* was highly expressed in stems and roots, followed by leaves(Fig. 4). The gene expression levels showed similar trends when they were normalized using the stable reference genes that we identified. However, normalizing using *H3* resulted in quite different results.

**Fig. 4.**
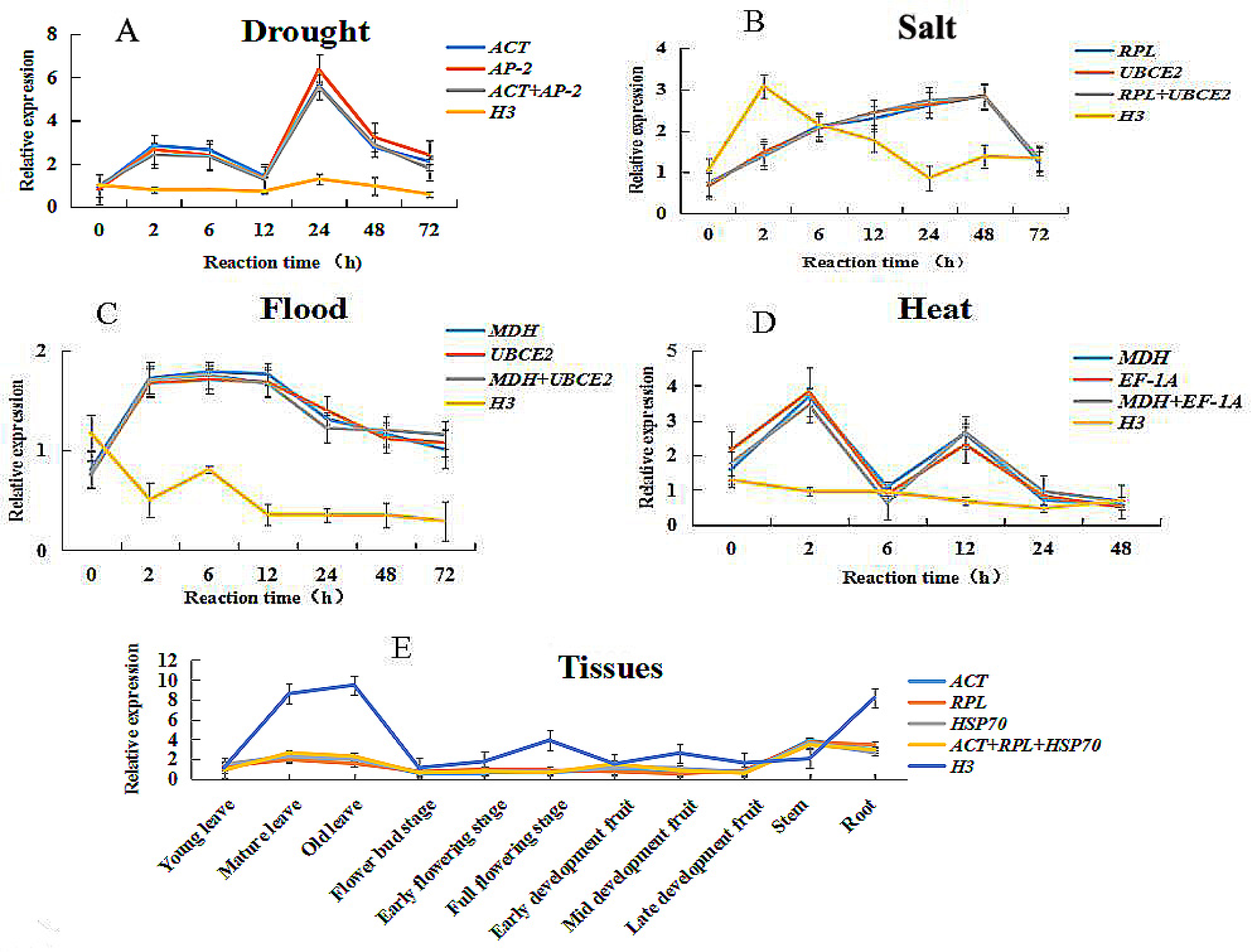
Relative quantification of *ClNHX1* and *ClLAC* expression using the validated reference genes. Leaves were collected from seedlings subjected to PEG, salt and flood stress after 0, 2, 6, 12, 24, 48 and 72 h (A, B, C). Leaves were collected from seedlings subjected to heat stress after 0, 2, 6, 12, 24 and 48 h (D). All abiotic stress treatments used the *ClNHX1* gene. Relative quantification of *ClLAC* expression using the validated reference genes in different tissues (E)

## Disscussion

Recently, qRT-PCR has been one of the most important techniques for detecting nucleic acid content and analyzing gene relative expression levels (Johnson et al., 2013). It has been widely used for high sensitivity, strong specificity and repeatable accuracy (Huggett et al., 2005; Derveaux et al., 2010). However, amplification products and RNA integrity of different lengths also affect expression analysis. In order to minimize the interference of RNA integrity on the results and the precise comparison of mRNA transcription in different samples, it is important to select the appropriate reference genes (Radonić et al., 2004; Obrero A et al., 2006). Traditional reference genes cannot be expressed stably in all cases, and their transcriptional stability is also species-dependent (Dheda et al., 2005). Therefore, in some cases, it is necessary to select the corresponding reference gene expressed at a constant level (Schmittgen et al., 2000; Tang et al., 2017). We designed primers of some traditional reference genes such as *UBQ* and *GAPDH* and of some new reference genes like *TIP41* and *DNAJ* (Shivhare et al., 2016; Zhang et al., 2017). However, our results showed that some traditional reference genes did not have a single specificity in *C. trichotomum*, which indicated that appropriate internal reference genes differed among species, experimental conditions and tissues.

*Cl.trichotomum* is a traditional medicinal plant and has a long flowering period with excellent ornamental characteristics. It also has excellent tolerance to various abiotic stresses, especially salt stress. Soil salinization is a worldwide problem (He et al., 2014) and coastal saline-alkali plant selection has been in the spotlight of coastal ecological restoration in recent decades (Ravindran et al., 2007; Li et al., 2010). Using qRT-PCR to select a stable reference gene as the normalization factor would ensure reliable qRT-PCR data for target genes and help us better understand the salt tolerance of plants at a molecular level. Here, 17 candidate genes were selected in *C. trichotomum*, and their expression stability in different tissues and various treatments were analyzed. Generally, three or more validated reference genes is considered to be appropriate for analyzing expression of target genes (Bustin et al., 2005). In this research, under various single abiotic stress treatments (drought, flood, salt, and heat) in TA and YA sources, the GeNorm results indicated that we should choose two candidate genes, while in tissues, all abiotic stresses and all samples, the results indicated that more than three genes were needed. This may be due to significant differences in gene expression of sources we selected with distant phylogenetic relationships. For convenience, when using internal reference genes, only the top three genes in the overall ranking should be chosen. Due to different statistical algorithms and analysis procedures, the application of different analysis software in the same tissue or treatment will lead to different verification results (Li et al., 2016). In our study, different software gave different results, but the stable ranking of genes was roughly the same.

Genes *UBCE2*, *RPL* and *ACT* have been demonstrated to be stable under abiotic stresses across various plants (Cameron et al., 2013; Shivhare et al., 2016; Li et al., 2017). In *Hibiscus cannabinus* and *Salicornia europaea*, *ACT* was also the optimum reference gene under drought stress (Niu et al., 2017). After expression analysis of candidate reference genes in *Urochloa brizantha*, *EF-1A* was demonstrated as the most suitable reference gene for heat stress (Stephan et al., 2019; Takamori et al., 2017), and *EF-1A* was also suitable in various plants (Narancio et al., 2011; Yang et al., 2014; Zhong et al., 2018). Gene *MDH* was confirmed to be the best reference gene in amaranth exposed to biotic or abiotic stress (Vera Hernandez et al., 2018), and was stable in various tissues of apple (Perini et al., 2014). Gene *AP-2* was identified as a potential reference gene due to stability in *Arabidopsis* and grapevine (Czechowski et al., 2005; Tashiro et al., 2016). Our results indicated that a combination of *RPL*, *ACT* and *HSP70* were the most suitable reference genes for tissues; in other plants, *RPL* and *ACT* were also the best reference genes for different tissues (Chen et al., 2011; Bin et al., 2012; You et al., 2018), and *HSP70* was identified as the candidate reference gene in carrot (Campos et al., 2014). Across all abiotic stresses, *ACT* and *AP-2* were stably expressed under drought stress, *MDH* and *UBCE2* under flood stress, and *UBCE2* and *RPL* under salt stress. For heat stress, *MDH* and *EF-1A* were the most suitable reference genes; and *RPL*, *MDH* and *AP-2* were the most appropriate under all four abiotic stresses. In general, *AP-2*, *RPL* and *ACT* were expressed stably in all samples. Gene *H3* was found to be the unstablereference gene in liverwort *Marchantia polymorpha* (Saint-Marcoux et al., 2015), and in our studies *H3* was also one of the most unstable reference gene. To verify the reliability of the reference genes we selected, the expression levels of *ClNHX1* and *ClLAC* were normalized by the identified reference genes, and results showed that the reference genes were stable in various abiotic stress treatments and tissues and indicated that different experimental conditions required different reference genes. This work provided guidance on the selection of reference genes for the application of *Clerodendrum trichotomum*, as well as reference gene selection for other *Clerodendrum* species. And this work will help to establish a better standardization and quantification of transcript levels in *C. trichotomum* in the future.

## Supporting information

**S1 Amplified nucleotide sequences.**

## Acknowledgments

This work was funded by the Project of National Forestry Public Welfare Industry Research Project (201404109), National Natural Science Foundation of China (Grant No. 31870695 and 31601785), the Top-notch Academic Programs Project of Jiangsu Higher Education Institutions and the Postgraduate Research & Practice Innovation Program of Jiangsu Province.

## Author Contributions

XY, YY, YH designed the experiment; YH conducted the experiment, analyzed the data and wrote the manuscript; and YH, GC,TY, WD, TS, LW and DH contributed to manuscript revision; and all authors read and approved the final manuscript.

## References

Andersen C L, Jensen J L, Orntoft T F. Normalization of real-time quantitative reverse transcription-PCR data: a model-based variance estimation approach to identify genes suited for normalization, applied to bladder and colon cancer data sets. Cancer Res. 2004; 64(15), 5245–5250.

Bin W S, Wei L K, Ping D W, Li Z, Wei G, Bing L J, et al. Evaluation of appropriate reference genes for gene expression studies in pepper by quantitative real-time PCR. Mol Breeding. 2012; 30(3), 1393–1400.

Borges A F, Fonseca C, Ferreira R B, Lourenco A M, Monteiro S. Reference gene validation for quantitative RT-PCR during biotic and abiotic stresses in Vitis vinifera. PLoS ONE. 2014; 9(10), e111399.

Stephan L, Tilmes V, Hülskamp M. Selection and validation of reference genes for quantitative Real-Time PCR in *Arabis alpina*. PLoS ONE. 2019;14(3): e0211172.

Bustin S A. Real-Time Reverse Transcription PCR. Encyclopedia of Genetics Genomics Proteomics & Informatics. 2005; 29(1), 1459.

Cameron R C, Duncan E J, Dearden P K. Stable reference genes for the measurement of transcript abundance during larval caste development in the honeybee. Apidologie. 2013; 44(4), 357–366.

Campos M D, Frederico A M, Nothnagel T, Arnholdt-Schmitt B, Cardoso H. Selection of suitable reference genes for reverse transcription quantitative real-time PCR studies on different experimental systems from carrot (Daucus carota L.). Sci Hortic-Amsterdam. 2015; 186, 115–123.

Chen L, Zhong H, Kuang J, Li J, Lu W, Chen J. Validation of reference genes for RT-qPCR studies of gene expression in banana fruit under different experimental conditions. Planta. 2011; 234(2), 377–390.

Czechowski T, Stitt M, Altmann T, Udvardi M K, Scheible W R. Genome-wide identification and testing of superior reference genes for transcript normalization in Arabidopsis. Plant Physiol. 2005; 139(1), 5–17.

Derveaux S, Vandesompele J, Hellemans J. How to do successful gene expression analysis using real-time PCR. Methods. 2010; 50(4), 227–230.

Dheda K, Huggett J F, Chang J S, Kim L U, Bustin S A, Johnson M A, et al. The implications of using an inappropriate reference gene for real-time reverse transcription PCR data normalization. Anal Biochem. 2005; 344(1), 141–143.

Fernandez P, Di Rienzo J A, Moschen S, Dosio G A, Aguirrezabal L A, Heinz R A. Comparison of predictive methods and biological validation for qPCR reference genes in sunflower leaf senescence transcript analysis. Plant Cell Rep. 2011; 30(1), 63–74.

Galli V, Borowski J M, Perin E C, Messias R S, Labonde J, Pereira I S, et al. Validation of reference genes for accurate normalization of gene expression for real time-quantitative PCR in strawberry fruits using different cultivars and osmotic stresses. Gene. 2015; 554(2), 205–214.

Gines M, Baldwin T, Bregitzer P, Maughan P J, Klos K E, Esvelt K. Selection of Expression Reference Genes with Demonstrated Stability in Barley among a Diverse Set of Tissues and Cultivars. Crop Sci. 2018; 58(1), 332–341.

He B, Cai Y, Ran W, Jiang H. Spatial and seasonal variations of soil salinity following vegetation restoration in coastal saline land in eastern China. Catena. 2014; 118, 147–153.

Hoagland D R, Arnon D I. The water-culture method for growing plants without soil. Calif Agric Exp Stn circ. 1950; 347(5406), 357–359.

Hu R, Fan C, Li H, Zhang Q, Fu Y. Evaluation of putative reference genes for gene expression normalization in soybean by quantitative real-time RT-PCR. BMC Mol Biol. 2009; 10(93).

Huggett J, Dheda K, Bustin S, Zumla A. Real-time RT-PCR normalisation; strategies and considerations. Genes Immun. 2005; 6(4), 279–284.

Iwatsuki K. Flora of Japan. Tokyo: Kodansha.1993.

Jha A, Joshi M, Yadav N S, Agarwal P K, Jha B. Cloning and characterization of the Salicornia brachiata Na+/H+ antiporter gene *SbNHX1* and its expression by abiotic stress. Mol Biol Rep. 2011; 38(3), 1965–1973.

Jiang Q, Wang F, Li M, Ma J, Tan G, Xiong A. Selection of Suitable Reference Genes for qPCR Normalization under Abiotic Stresses in *Oenanthe javanica* (BI.) DC. PLoS ONE. 2014; 9(e922623).

Johnson G, Nolan T, Bustin S A. Real-time quantitative PCR, pathogen detection and MIQE. Methods in molecular biology (Clifton, N.J.). 2013; 943, 1–16.

Kitajima S, Imamura T, Iibushi J, Ikenaga M, Tachibana Y, Andoh N, et al. Ferritin 2 domain-containing protein found in lacquer tree (*Toxicodendron vernicifluum*) sap has negative effects on laccase and peroxidase reactions. Biosci Biotech Bioch. 2017; 81(6), 1165–1175.

Li M, Wang F, Jiang Q, Wang G, Tian C, Xiong A. Validation and Comparison of Reference Genes for qPCR Normalization of Celery (*Apium graveolens*) at Different Development Stages. Front Plant Sci. 2016; 7, 313.

Li T, Wang J, Lu M, Zhang T, Qu X, Wang Z. Selection and Validation of Appropriate Reference Genes for qRT-PCR Analysis in *Isatis indigotica* Fort. Front Plant Sci. 2017; 8, 1139.

Li Z, Li G, Qin P. The prediction of ecological potential for developing salt-tolerant oil plants on coastal saline land in *Sheyang Saltern*, China. Ecol Eng. 2010; 36(1), 27–35.

Li Z, Zhang J, Qn L, Ge Y. Enhancing Antioxidant Performance of Lignin by Enzymatic Treatment with Laccase. Acs Sustain Chem Eng. 2018; 6(2), 2591–2595.

Lilly S T, Drummond R S M, Pearson M N, MacDiarmid R M. Identification and Validation of Reference Genes for Normalization of Transcripts from Virus-Infected *Arabidopsis thaliana*. Mol Plant Microbe In. 2011; 24(3), 294–304.

Machado R D, Christoff A P, Loss-Morais G, Margis-Pinheiro M, Margis R, Korbes A P. Comprehensive selection of reference genes for quantitative gene expression analysis during seed development in *Brassica napus*. Plant Cell Rep 2015; 34(7), 1139–1149.

Mahmud R, Inoue NKasajima S Y, Shaheen R. Assessment of potential indigenous plant species for the phytoremediation of arsenic-contaminated areas of Bangladesh. Int J Phytoremediat. 2008; 10(2), 119–132.

Mamidala P, Rajarapu S P, Jones S C, Mittapalli O. Identification and validation of reference genes for quantitative real-time polymerase chain reaction in *Cimex lectularius*. J Med Entomol. 2011; 48(4), 947–951

Mu H, Sun T, Xu C, Wang L. Yue Y, Yang X. Identification and validation of reference genes for gene expression studies in sweet osmanthus (*Osmanthus fragrans*) based on transcriptomic sequence data. J Genet. 2017; 96(2), 273–281.

Narancio R, John U, Mason J, Spangenberg G. Selection of optimal reference genes for quantitative RT-PCR transcript abundance analysis in white clover (*Trifolium repens* L.). Funct Plant Biol. 2018; 45(7), 737–744.

Niu X, Chen M, Huang X, Chen H, Tao A, Qi J. Reference Gene Selection for qRT-PCR Normalization Analysis in kenaf (*Hibiscus cannabinus* L.) under Abiotic Stress and Hormonal Stimuli. Front Plant Sci. 2017; 8, 771.

Obrero A, Die J V, Román B, Gómez P, Nadal S, González-Verdejo CI. Selection of reference genes for gene expression studies in zucchini (*cucurbita pepo*) using qpcr. J Agric Food Chem. 2011; 59(10), 5402–5411.

Perini P, Pasquali G, Margis-Pinheiro M, Dias De Oliviera P R, Revers L F. Reference genes for transcriptional analysis of flowering and fruit ripening stages in apple (*Malus × domestica* Borkh.). Mol Breeding. 2014; 34(3), 829–842.

Pfaffl M W. A new mathematical model for relative quantification in real-time RT-PCR. Nucleic Acids Res. 2001; 29(9), e45

Pfaffl M W, Tichopad A, Prgomet C, Neuvians T P. Determination of stable housekeeping genes, differentially regulated target genes and sample integrity: BestKeeper--Excel-based tool using pair-wise correlations. Biotechnol Lett; 2004. 26(6), 509–515

Qi S, Yang L, Wen X, Hong Y, Song X, Zhang M, et al. Reference Gene Selection for RT-qPCR Analysis of Flower Development in *Chrysanthemum morifolium* and *Chrysanthemum lavandulifolium*. Front Plant Sci. 2016; 7, 287.

Radonic A, Thulke S, Mackay I M, Landt O, Siegert W, Nitsch A. Guideline to reference gene selection for quantitative real-time PCR. Biochem Bioph Res Co. 2004; 313(4), 856–862.

Ravindran K C, Venkatesan K, Balakrishnan V, Chellappan K P, Balasubramanian T. Restoration of saline land by halophytes for Indian soils. Soil Biol Biochem. 2007; 39(10), 2661–2664.

Rodriguez H A, Rodriguezarango E, Morales J G, Kema G H J, Arango R. Defense Gene Expression Associated with Biotrophic Phase of Mycosphaerella fijiensis M. Morelet Infection in Banana. Plant Dis. 2016; 100(6), 1170–1175.

Sahoo D P, Kumar S, Mishra S, Kobayashi Y, Panda S K, Sahoo L. Enhanced salinity tolerance in transgenic mungbean overexpressing Arabidopsis antiporter (*NHX1*) gene. Mol Breeding. 2016; 36(14410).

Saint-Marcoux D, Proust H, Dolan L, Langdale J A. Identification of reference genes for real-time quantitative PCR experiments in the liverwort Marchantia polymorpha. PLoS ONE. 2015; 10(3), e118678.

Savard P, Roy D. Determination of Differentially Expressed Genes Involved in Arabinoxylan Degradation byBifidobacterium longumNCC2705 Using Real-Time RT-PCR. Probiotics Antimicro. 2009; 1(2), 121–129.

Schmittgen Z, Schmittgen T D, Zakrajsek B A. Effect of experimental treatment on housekeeping gene expression: validation by real-time, quantitative RT-PCR. J Biochem Biophys Methods. 2000; 46, 69–81.

Shivhare R, Lata C. Selection of suitable reference genes for assessing gene expression in pearl millet under different abiotic stresses and their combinations. 2016; Sci. Rep. 6(23036).

Takamori L M, Pereira A, Maia S G, Vieira L, Ferreira R A. Identification of Endogenous Reference Genes for RT-qPCR Expression Analysis in *Urochloa brizantha* Under Abiotic Stresses. Sci Rep. 2017; 7(1), 8502.

Tang X, Zhang N, Si H, Calderon-Urrea A. Selection and validation of reference genes for RT-qPCR analysis in potato under abiotic stress. Plant Methods. 2017; 13(85).

Tashiro R M, Philips J G, Winefield C S. Identification of suitable grapevine reference genes for qRT-PCR derived from heterologous species. Mol Genet Genomics. 2016; 291(1), 483–492.

Tian C, Jiang Q, Wang F, Wang G, Xu Z, Xiong A. Selection of Suitable Reference Genes for qPCR Normalization under Abiotic Stresses and Hormone Stimuli in Carrot Leaves. PLoS ONE. 2015; 10(e01175692).

Vandesompele J, De Preter K, Pattyn F, Poppe B, Van Roy N, De Paepe A, et al. Accurate normalization of real-time quantitative RT-PCR data by geometric averaging of multiple internal control genes. Genome Biol. 2002; 3(7), H34.

Vera Hernandez F P, Martinez Nunez M, Ruiz Rivas M, Vazquez Portillo R E, Bibbins Martinez M D, Rosas Cardenas F D F. Reference genes for RT-qPCR normalisation in different tissues, developmental stages and stress conditions of amaranth. Plant Biology. 2018; 20(4), 713–721.

Wang J, Luan F, He X, Wang Y, Li M. Traditional uses and pharmacological properties of Clerodendrum phytochemicals. J Altern Complem Med. 2018; 8(1), 24–38.

Wang T, Hao R, Pan H, Cheng T, Zhang Q. Selection of Suitable Reference Genes for Quantitative Real-time Polymerase Chain Reaction in *Prunus mume* during Flowering Stages and under Different Abiotic Stress Conditions. J Am Soc Hortic Sci. 2014; 139(2), 113–122

Wang W X, Xiong J, Tang Y, Zhu J J, Li M, Zhao Y, et al. Rearranged abietane diterpenoids from the roots of *Clerodendrum trichotomum* and their cytotoxicities against human tumor cells. Phytochemistry. 2013a; 89, 89–95.

Wang W, Zhu J, Zou Y, Hong Z, Liu S, Li M, et al. Trichotomone, a new cytotoxic dimeric abietane-derived diterpene from *Clerodendrum trichotomum*. Tetrahedron Lett. 2013b; 54(20), 2549–2552.

Xu H, Li J, Wu R, Su W, Wu X, Wu X, et al. Identification of Reference Genes for Studying Herbicide Resistance Mechanisms in Japanese Foxtail (*Alopecurus japonicus*). Weed Sci. 2017, 65(5), 557–566.

Xu R L, Wang R, Wei H, Shi Y P. New cyclohexylethanoids from the leaves of *Clerodendrum trichotomum*. Phytochem Lett. 2014; 7(2), 111–113.

Xu Y, Li H, Li X, Lin J, Wang Z, Yang Q, et al. Systematic selection and validation of appropriate reference genes for gene expression studies by quantitative real-time PCR in pear. Acta Physiol Plant. 2015; 37(UNSP 402).

Yang X, Li H, Yue Y, Ding W, Xu C, Wang L. Transcriptomic Analysis of the Candidate Genes Related to Aroma Formation in Osmanthus fragrans. Molecules. 2018; 23(16047).

Yang Q, Yin J, Li G, Qi L, Yang F, Li G. Reference gene selection for qRT-PCR in Caragana korshinskii Kom. under different stress conditions. Mol Biol Rep. 2014; 41(4), 2325–2334.

Yang Z, Chen Y, Hu B, Tan Z, Huang B. Identification and validation of reference genes for quantification of target gene expression with quantitative real-time PCR for tall fescue under four abiotic stresses. PLoS ONE. 2015; 10(3), e119569.

You Y, Xie M, Vasseur L, You M. Selecting and validating reference genes for quantitative real-time pcr in *plutella xylostella* (l.). Genome. 2018; gen-2017-0176.

Yue Y, Tian S, Wang Y, Ma H, Liu S, Hu H. Transcriptomic and GC-MS Metabolomic Analyses Reveal the Sink Strength Changes during Petunia Anther Development. Int J Mol Sci. 2018; 19(9554).

Yue Y, Yin C, Guo R, Peng H, Yang Z, Hu H. An anther-specific gene PhGRP is regulated by *PhMYC2* and causes male sterility when overexpressed in petunia anthers. Plant Cell Rep. 2017; 36(9), 1401–1415.

Zhang K, Li M, Cao S, Sun Y, Long R, Kang J, et al. Selection and validation of reference genes for target gene analysis with quantitative real-time PCR in the leaves and roots of *Carex rigescens* under abiotic stress. Ecotox Environ Safe 2019; 168, 127–137.

Zhang Y, Han X, Chen S, Zheng L, He X, Liu M, et al. Selection of suitable reference genes for quantitative real-time PCR gene expression analysis in *Salix matsudana* under different abiotic stresses. Sci Rep. 2017; 7, 40290.

Zhong H Y, Chen J W, Li C Q, Chen L, Wu J Y, Chen J Y, et al. Selection of reliable reference genes for expression studies by reverse transcription quantitative real-time PCR in litchi under different experimental conditions. Plant Cell Rep 2011; 30(4), 641–653.

